# Modeling diverse genetic subtypes of lung adenocarcinoma with a next-generation alveolar type 2 organoid platform

**DOI:** 10.1101/2021.12.07.471632

**Authors:** Santiago Naranjo, Christina M. Cabana, Lindsay M. LaFave, Peter M.K. Westcott, Rodrigo Romero, Arkopravo Ghosh, Laura Z. Liao, Jason M. Schenkel, Isabella Del Priore, Arjun Bhutkar, Dian Yang, Tyler Jacks

## Abstract

Lung cancer is the leading cause of cancer-related death worldwide. Lung adenocarcinoma (LUAD), the most common histological subtype, accounts for 40% of all cases. While genetically engineered mouse models (GEMMs) recapitulate the histological progression and transcriptional evolution of human LUAD, they are slow and technically demanding. In contrast, cell line transplant models are fast and flexible, but are often derived from clonal idiosyncratic tumors that fail to capture the full spectrum of clinical disease. Organoid technologies provide a means to create next-generation cancer models that integrate the most relevant features of autochthonous and transplant-based systems, yet robust and faithful LUAD organoid platforms are currently lacking. Here, we describe optimized conditions to continuously expand murine alveolar type 2 cells (AT2), a prominent cell-of-origin for LUAD, in organoid culture. These organoids display canonical features of AT2 cells, including marker gene expression, the presence of lamellar bodies, and an ability to differentiate into the AT1 lineage. We used this system to develop flexible and versatile immunocompetent organoid-based models of *KRAS* and *ALK-*mutant LUAD. Notably, the resultant tumors closely resemble their autochthonous murine counterparts and human LUAD. In contrast to comparable organoid platforms, our system supports long-term maintenance of the AT2 cellular identity, providing unprecedented ease and reliability to study AT2 and LUAD biology *in vitro* and *in vivo*.

## MAIN TEXT

We and others have developed various GEMMs to study LUAD^1–3^. In these models, tumors arise from lung epithelial cells and follow a progression pattern that faithfully recapitulates their human counterparts at the molecular and histopathological level. While GEMMs offer a physiologically relevant platform to study LUAD, they are time-consuming and technically demanding. On the other hand, traditional cancer cell line-based transplant models are genetically tractable and generate fast-growing tumors, but have been shown to acquire confounding genomic and transcriptomic alterations in culture^4^.

Organoid technology has provided a powerful means to create improved cancer models that are both practical and faithfully recapitulate clinical disease. Organoids are cultured mini organs derived from healthy stem cells that proliferate continuously while retaining their physiological identity^5–8^. Importantly, organoid transplant models of colorectal and pancreatic cancer closely resemble clinical disesase^9,10^. Altogether, organoids allow for the creation of models that are easy to manipulate experimentally and closely mimic the natural process of tumor initiation and progression.

AT2 cells are epithelial stem cells in the lung and represent a major cell-of-origin for LUAD^11–13^. AT2 organoid culture systems to date have been limited by significant technical and biological challenges. First generation systems require co-culture with feeder cells or an air-liquid-interface (ALI), making them experimentally inflexible and financially demanding^14^. Moreover, organoids cultured in this manner have been shown to differentiate into club^15^ or basal cell lineages^16^. While some of these challenges have been recently addressed by systems that use chemically-defined media without feeder cells or an ALI^17–20^, these platforms continue to display significant limitations. For example, it remains unclear whether one of these systems supports passaging^17^. Others display lengthy expansion periods between passages (1-2 months)^18,19^, with loss of AT2 identity over time^19^. These limitations have prevented significant advancements in modeling LUAD with AT2 organoids.

To our knowledge, only one group has described an orthotopic organoid transplant model of LUAD, which has two major limitations^21^. Introduction of oncogenic mutations into first generation AT2 organoids immediately induced transcriptional and morphological changes consistent with the profile of late stage LUAD^22^. Therefore, this system precludes the study of earlier stages of this disease. Furthermore, tumor penetrance and disease burden were not explored, and it remains unclear if these organoids can form tumors in immunocompetent mice.

Here, we describe an optimized AT2 organoid platform that employs chemically-defined media, circumvents feeder cells and ALI systems, and can be used to faithfully model LUAD *in vivo*. We demonstrate that these organoids can be expanded continuously and maintenance of the canonical, anatomical and transcriptional characteristics of AT2 cells. Importantly, we show that activating mutations in either *Kras* or *Alk* in combination with *Trp53* (*TP53* in humans) loss – three common driving events in human LUAD — permitted organoid proliferation in the absence of growth factors. These *in vitro* transformed organoids form tumors that display multi-stage progression in immunocompetent hosts that recapitulate key features of human LUAD.

## RESULTS

To build organoid-based models of LUAD, we undertook a systematic approach to define a cocktail of growth factors that could expand adult murine AT2 cells embedded in matrigel and submerged in medium. We hypothesized that combining activators of alveolar regeneration and proliferation (Fgfr2^23^, c-Met^24^, and Wnt^25,26^) with inhibitors of pathways that antagonize these processes (Bmp^27^, Tgf-ß^28^, and p38 MAPK^29^) would accomplish this goal.

To test this hypothesis, we embedded freshly dissociated bulk lung cell suspensions from transgenic *Sftpc-eGFP* mice^30^ in matrigel droplets and overlaid F^7^NHCSA medium (<u>F</u>GF<u>7</u>, <u>N</u>OGGIN, <u>H</u>GF, <u>C</u>HIR99021, <u>S</u>B202190 and <u>A</u>83-01) (**Supplementary Fig. 1A**). The *Sftpc-eGFP* transgene fluorescently labels cells expressing the canonical AT2 marker gene, *Sftpc*. Although we initially observed robust expansion of eGFP+ organoids, indicating maintenance of the AT2 state, eGFP-organoids progressively took over the cultures (**Supplementary Fig. 1B**). To transcriptionally define both populations, we performed bulk RNA sequencing (RNA-seq) on sorted eGFP+ and eGFP-cells. Gene set enrichment analysis (GSEA)^31,32^ revealed that genes from an established AT2 transcriptional signature^33^ were highly enriched in the eGFP+ group (**Supplementary Fig. 1C**). On the other hand, a gene set associated with pulmonary basal cells^34^correlated strongly with the eGFP-group (**Supplementary Fig. 1C**). These results demonstrate that eGFP+ and eGFP-cells in our culture system likely represent AT2 and basal cells, respectively.

By analyzing these transcriptional data, we identified *MHC-II* and *Egfr* as candidate surface markers for the eGFP+ (AT2) and eGFP-(basal) populations, respectively (**Supplementary Fig. 1E**). We validated these expression patterns at the protein level by flow cytometry (**Supplementary Fig. 1F**). Given the specificity of EGFR and MHC-II for the basal and alveolar states, respectively, these surface proteins are valuable markers for rapid and quantitative assessment of AT2 cell state fidelity in lung organoid cultures.

Next, we tested a modified protocol to achieve expansion of AT2 organoids that retain their identity over time. We hypothesized that the presence of multiple cell types in the initial culture contributed to outgrowth of basal organoids. Thus, instead of bulk dissociated lung, we seeded freshly sorted AT2 cells as a starting population^35^.

Furthermore, we used F^7^NHCS medium (**Fig. 1A**), which lacks A-83-01, as early experiments suggested this factor exacerbated the growth of eGFP-organoids. This optimized protocol supported long-term expansion (at least 8 passages) of AT2 organoids in 8 out of 10 lines tested (**Fig. 1B-D**). Importantly, lines that maintained alveolar identity throughout the first two passages remained stable long-term (**Fig. 1D**). Lastly, we derived organoids from three mice lacking the *Sftpc-eGFP* reporter and demonstrated that they remained positive for MHCII and negative for EGFR for at least 6 passages (**Supplementary Fig. 1G**). Collectively, these results demonstrate that our optimized protocol robustly expands AT2 cells that retain their identity over time.

**Figure 1.**
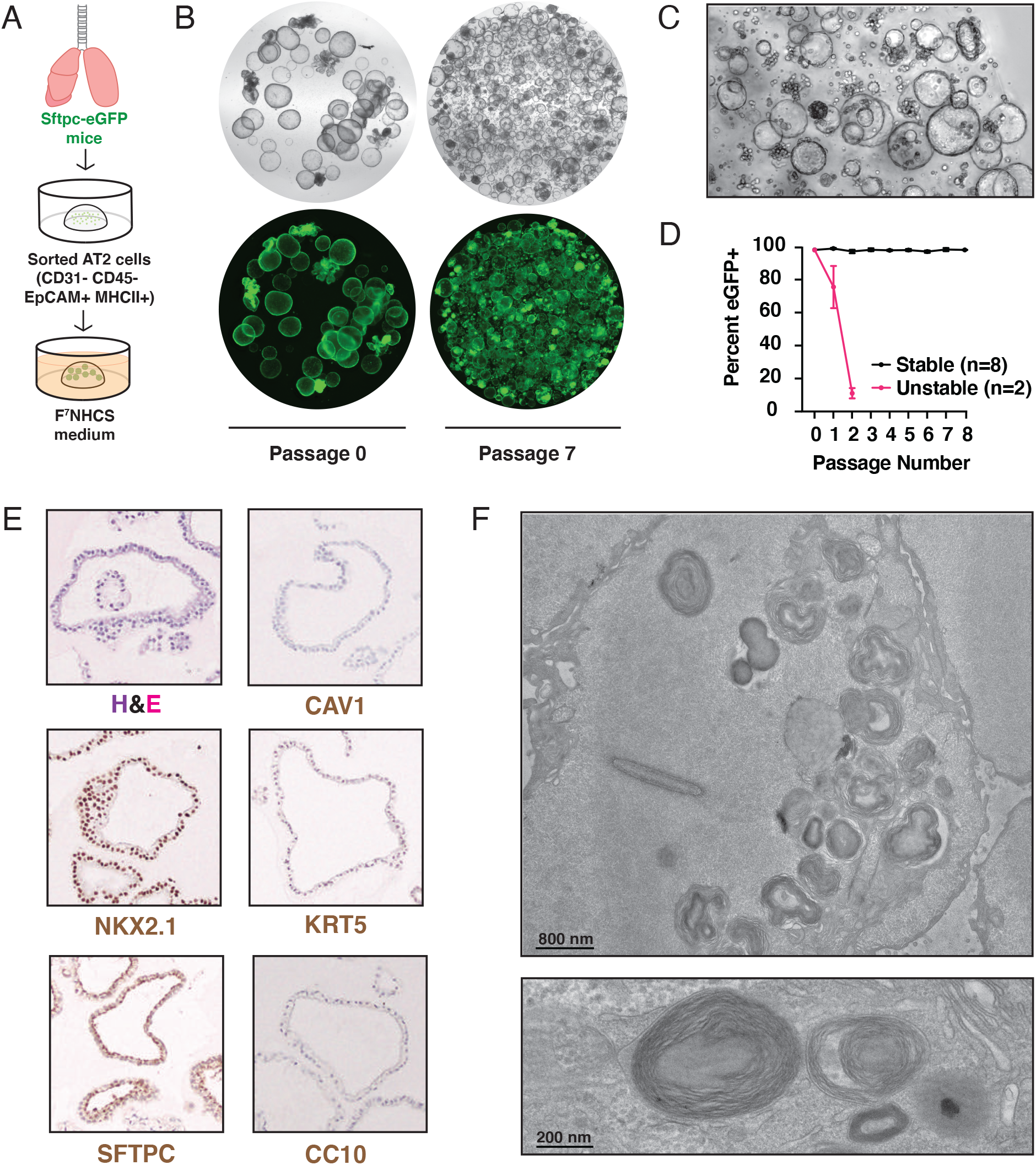
A. Method to culture alveolar organoids from normal lungs. B. Representative images of Sftpc-eGFP organoids at an early (0) and late (7) passage. C. Representative brightfield image of AT2 organoids, 10x magnification. D. Flow cytometry-based quantification of Sftpc-eGFP expressing cells over time in organoid culture. A total of *n* = 10 independent lines were established from Sftpc-eGFP mice and tested over three different experiments. All data are expressed as mean values ± the standard deviation. E. Hematoxylin and eosin (H&E) and immunohistochemical staining of organoids for canonical lung cell markers including SFTPC (AT2 cells), CAV1 (AT1 cells), KRT5 (Basal cells), CGRP (Pulmonary neuroendocrine cells), CC10 (Club cells). F. Electron microscopy images of organoids showing lamellar body structures.

Consistent with maintenance of the AT2 state, these organoids expressed SFTPC as indicated by immunohistochemical staining, but lacked expression of markers of other major pulmonary epithelial cell types (**Fig. 1E**). Likewise, the cells comprising these organoids were densely packed with lamellar bodies, an organelle that stores and secretes pulmonary surfactant in primary AT2 cells (**Fig. 1F**).

We next tested the ability of these organoids to differentiate into an AT1 state – a characteristic feature of AT2 cells – *in vitro* and *in vivo*. First, we compared the morphology of organoids cultured in normal (F^7^NHCS - 3D) culture conditions against two experimental monolayer conditions using either complete media (F^7^NHCS - 2D) or differentiation media (10%FBS - 2D) (**Fig. 2A**). Cells grown under F^7^NHCS - 2D conditions adopted a cuboidal morphology, consistent with what is typically observed for AT2 cells. In stark contrast, cells grown in 10%FBS - 2D conditions adopted an elongated and flat morphology that is characteristic of AT1 cells (**Fig. 2B**).

**Figure 2.**
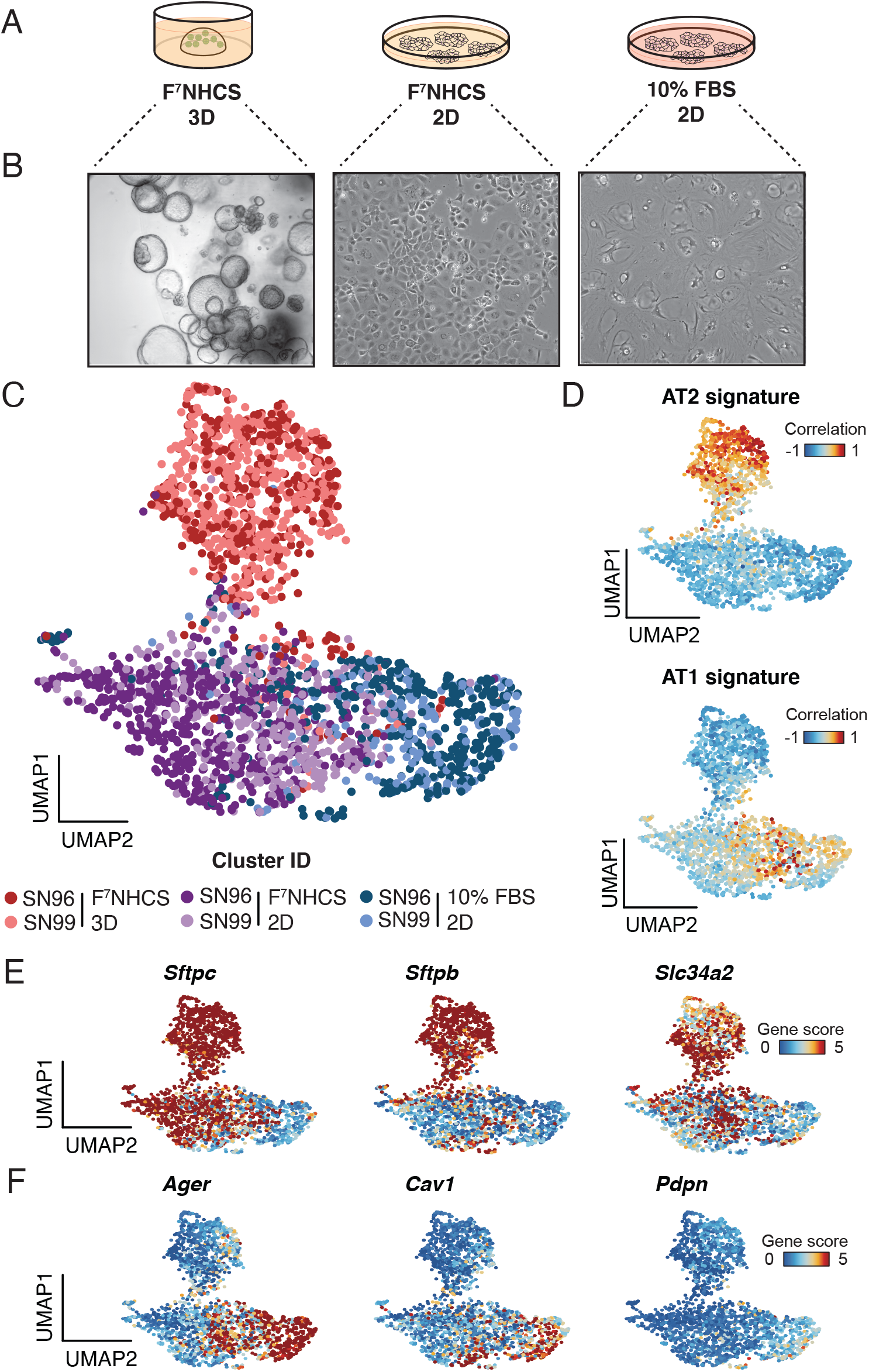
A. Schematic of culture conditions for *in vitro* differentiation experiment. B. Brightfield images of organoid cells after one week in specified culture conditions (10x, 20x, and 20x magnification, respectively). C. UMAP visualization of organoids (*n* = 2) grown in the indicated culture conditions and profiled by sci-ATAC-seq. Organoids grown in F^7^NHCS – 3D are colored in red, F^7^NHCS – 2D in purple, and 10% FBS – 2D in blue. D. Correlation of chromatin profiles in single organoid cells to established chromatin accessibility signatures for AT2 and AT1 cells. Cells are colored by their Pearson’s differential correlation coefficients. E. UMAP highlighting single-cell gene scores for three canonical AT2 (*Lamp3, Lyz2*, and *Sftpc*) markers in organoids cultured in the three conditions indicated in *Fig. 2A*. F. UMAP highlighting single-cell gene scores for three canonical AT1 (*Ager, Pdpn*, and *Cav1*) markers in organoids cultured in the three conditions indicated in *Fig. 2A*.

To test whether our experimental conditions induced full or partial differentiation to the AT1 lineage at the molecular level, we performed <u>s</u>ingle cell <u>c</u>ombinatorial indexing <u>a</u>ssay for transposase-<u>a</u>ccessible <u>c</u>hromatin <u>seq</u>uencing (sci-ATAC-seq)^36^, which measures genome-wide chromatin accessibility in single cells across a population. We visualized these data using <u>U</u>niform <u>M</u>anifold <u>A</u>pproximation and <u>P</u>rojection (UMAP)^37^. Our analysis showed that cells in the F^7^NHCS - 3D condition display unique chromatin accessibility profiles relative to those in the other two conditions, which overlapped significantly (**Fig. 2C**). Cells in the F^7^NHCS - 3D condition had high chromatin accessibility at genes comprising an epigenomic signature derived from primary AT2 cells^38^ (**Fig. 2D, Fig. 2E**), while cells in the F^7^NHCS - 2D and 10% FBS – 2D groups were moderately and highly concordant with a primary AT1 epigenomic signature^38^, respectively (**Fig. 2D, Fig. 2F**).

The above results led us to hypothesize that the 2D culture environment pushes the cells to partially differentiate into the AT1 state, while the F^7^NHCS medium promotes retention of key features of AT2 cells. This hypothesis is supported by the heterogenous accessibility at the individual AT2 maker loci (**Fig. 2E**) for cells in the F^7^NHCS - 2D condition. In contrast, the 10% FBS supplemented medium induced full differentiation into the AT1 state (**Fig. 2D, Fig. 2F**). Importantly, none of these conditions promoted chromatin accessibility profiles that correlated significantly with signatures of other major lung epithelial cell types (**Supplementary Fig. 2A**). We next applied bulk RNA-seq to examine whether the transcriptional states of these organoids were consistent with their epigenomic profiles. As expected, GSEA confirmed that the F^7^NHCS - 3D condition maintained the AT2 state, while the 10% FBS - 2D condition induced an AT1 state (**Supplementary Fig. 2B**). Again, neither sample correlated with club nor basal transcriptomic states (**Supplementary Fig. 2B**).

To test the ability of our organoids to self-renew and differentiate *in vivo*, we intratracheally transplanted wildtype organoids tagged with a lentivirus expressing eGFP driven by the EF1a promoter into syngeneic mice pre-treated with low-dose bleomycin, as this has been shown to enhance the engraftment efficiency of cells into the lung^39^. Engrafted eGFP+ cells were detected in the lung parenchyma of 5/7 transplanted mice, albeit at low numbers. Some engrafted cells maintained SFTPC expression (**Supplementary Fig. 2C**), while others upregulated CAV1 (**Supplementary Fig. 2D**). Importantly, eGFP+ cells did not stain positively for CD45, indicating that they had engrafted into the lung parenchyma and not phagocytosed by macrophages. These data suggest that organoid-derived AT2 cells retain the capacity to differentiate into AT1 cells *in vivo*.

We next utilized our optimized organoid system to model clinically-relevant LUAD subtypes. *KRAS* is mutated in ∼30% of LUAD patients while *ALK* fusions are observed in ∼1% of clinical cases^40^. On the other hand, *TP53* is mutated in ∼50% of patient lung adenocarcinomas^40^. Importantly, *KRAS*^*G12D*^ and *EML4*-*ALK* alone are able to initiate lung tumors in mice^3,41^. While *Trp53* loss is not sufficient for tumorigenesis, this genetic event accelerates LUAD progression in *Kras*- or *Braf*-driven GEMMs^2,42^. We therefore modeled *KRAS*^*G12D*^ or *EML4*-*ALK* alterations in combination with *TP53* loss.

To test whether these prevalent genetic insults could transform wildtype AT2 organoids, we created *<u>K</u>ras*-mutant, *Tr<u>p</u>53*-deficient (*KP)* organoids by infecting *Kras*^*LSL-G12D/+*^*;Trp53*^*fl/fl*^*;Rosa26*^*LSL-Cas9-eGFP/+*^ organoids with adenoviruses expressing Cre recombinase (*Ad5-Cre*). Likewise, we generated *<u>E</u>ml4*-*<u>A</u>lk*-mutant, *Tr<u>p</u>53*-deficient (*EAP)* organoids by transducing *Trp53*^*fl/fl*^*;Rosa26*^*LSL-Cas9-eGFP*^ organoids with an adenovirus expressing Cre and sgRNAs (*Ad5-U6-sgEml4-U6-sgAlk-CBh-Cre*), that were previously shown to induce a clinically-relevant oncogenic fusion between *Eml4* and *Alk* ^3^. As a control, we transduced *Trp53*^*fl/fl*^*;R26*^*LSL-Cas9-eGFP*^ organoids with *Ad5-Cre* to create *Tr<u>p</u>53*-deficient only organoids (*P only*) (**Fig. 3A**). In each case, the recombined *R26*^*LSL-Cas9-eGFP*^ allele provided a fluorescent tag that allowed us to easily visualize mutant organoids. PCR analysis demonstrated efficient recombination of the *Kras*^*LSL-G12D/+*^ allele and induction of *Eml4-Alk* inversion (**Supplementary Fig. 2E**).

**Figure 3.**
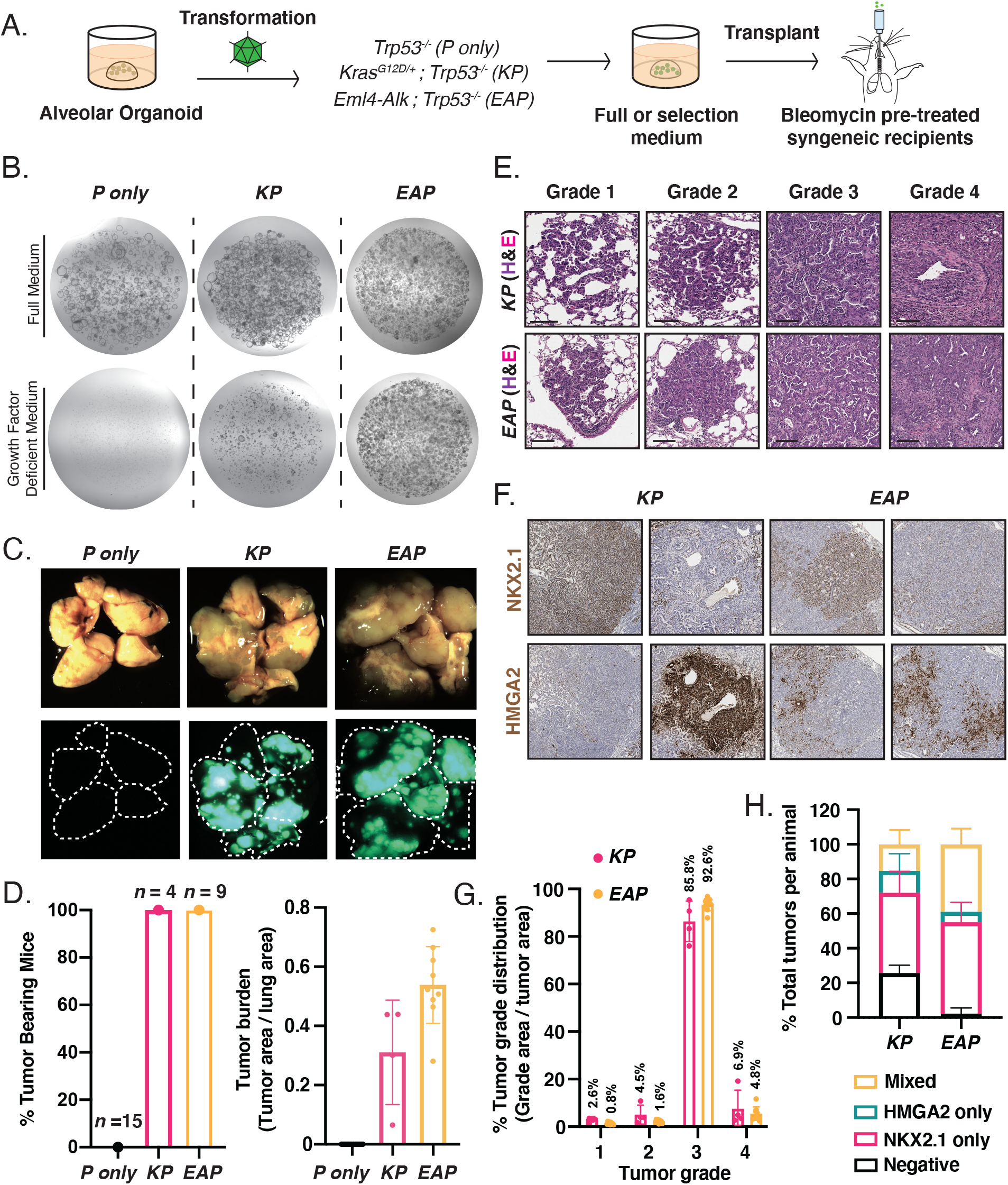
A. Strategy to introduce the indicated sets of oncogenic mutations (see main text for specific details) and test their effect on AT2 organoids *in vitro* and *in vivo*. B. Representative images of transformed (*KP* and *EAP*) and normal (*P only*) organoids in full or selection medium lacking growth factors. C. Representative gross brightfield (top) and fluorescent (bottom) images of lungs from recipient mice 4 months (*KP* and *P only)* and 2 months (*EAP*) post transplantation. D. Quantification of penetrance for tumor formation (left) and tumor burden (right). *P only* is shown in black, *KP* in pink, and *EAP* in yellow. E. Hematoxylin and eosin (H&E) staining of representative tumors across all grades from transplanted *KP* (top row) and *EAP* (bottom row) organoids. F. Immunohistochemical staining of organoid derived tumors for canonical markers of early (*Nkx2*.*1*) and late-stage (*Hmga2*) lung adenocarcinoma. G. Quantification of tumor burden by grade. *KP* shown in pink, and *EAP* in yellow. Average value listed above each bar. H. Quantification of tumor lesions which stained positive for NKX2.1, HMGA2, both or neither via immunohistochemical staining.

We subsequently cultured these lines in complete (F^7^NHCS) and growth-factor depleted media (CS, selection media) to assay for oncogenic transformation. *P* only, *KP*, and *EAP* organoids were able to expand in complete media without any obvious growth differences. In stark contrast, only the organoids carrying activated oncogenes were able to grow in media lacking growth factors and Noggin (CS media) (**Fig. 3B**). To test the tumorigenic potential of these lines *in vivo*, we orthotopically transplanted them into syngeneic, immunocompetent mice (**Fig. 3C**). Prior to transplantation, we intratracheally dosed mice with low-dose bleomycin^39^. Notably, *KP* and *EAP* organoids gave rise to macroscopic tumors in 100% of recipient mice and yielded extensive tumor burden (**Fig. 3D**).

Next, we probed the validity of our platform to recapitulate human LUAD. *KP* and *EAP* tumors displayed histopathological features that are characteristic of human LUAD and recapitulated the full spectrum of tumor progression, including hyperplasias (Grade 1), adenomas (Grade 2), adenocarcinomas (Grade 3), and invasive adenocarcinomas (Grade 4) (**Fig. 3E**). The majority of tumors across both genotypes were Grade 3, consistent with the late time-point at which the recipients were analyzed (*KP* ∼ 16 weeks, *EAP* ∼ 8 weeks; **Fig. 3G**). Importantly, late-stage tumors in the autochthonous *KP* model display a similar grade distribution, further supporting the validity of this organoid platform^43^.

*Nkx2*.*1* and *Hmga2* are robust markers of benign and malignant lung tumor progression stages, respectively, in the autochthonous KP model^22^. To assess tumor progression at a molecular level, we measured NKX2.1 and HMGA2 protein expression in these tumors. Most KP and EAP tumors (∼75% and ∼98%, respectively) expressed NKX2.1, HMGA2, or a mixture of both (**Fig. 3F, 3H**). Individual tumors that were classified as “mixed” predominately contained areas that were positive for one marker but not the other. These results further demonstrate that our organoid models recapitulate fundamental histopathological and molecular features of LUAD throughout multiple stages of progression.

## DISCUSSION

Organoids provide a powerful platform to model cancer, but the lack of stable AT2 organoid culture systems has precluded the application of this technology to develop next generation models of LUAD. Here, we designed and validated a chemically defined growth factor cocktail that supports sustained growth and expansion of murine organoids displaying cellular and molecular canonical features of AT2 cells. Furthermore, we demonstrate that AT2 organoids can be genetically manipulated using a combination of Cre/loxP and CRISPR/Cas9 technologies followed by selection and orthotopic transplantation to rapidly develop clinically-relevant organoid models of LUAD. As proof-of-concept, we deployed this system to rapidly develop LUAD models driven by *KP* and *EAP* genetic lesions and show that they produced tumors that are highly faithful to human disease and indistinguishable from their autochthonous ‘gold-standard’ counterparts.

Our organoid-based cancer models provide significant advantages over previous platforms. First, our system permits rapid interrogation of diverse genetic lesions. In this study, we illustrated rapid engineering of a chromosomal inversion that leads to an oncogenic fusion between *Eml4* and *Alk* using CRISPR/Cas9. While our proof-of-concept experiments employed Cre recombinase or the Cas9 nuclease, we anticipate that other genome editing technologies (such as Base^44^ or Prime^45^editing) or transcriptional modulation technologies (such as CRISPR-based gene activation^46^ or repression^47^) can be seamlessly deployed to engineer these organoids at any desired scale. This level of flexibility should accelerate efforts centered on functional characterization of genetic alterations that continue being identified in patient tumors through large-scale genomic^40^ and transcriptomic^48^ profiling studies. Likewise, combining genetic screens with genetically defined normal and transformed organoid pairs can help uncover tumor-specific and even genotype-specific vulnerabilities.

Moreover, our model employs immunocompetent hosts and yields extensive tumor burden with 100% penetrance. These two features combined will enable studies of immune dysfunction, emerging immunotherapies in lung cancer, and, potentially, the function of MHCII in LUAD, a topic that remains poorly understudied. Organoids could also be rendered immunogenic by the inclusion of model antigens^49,50^ or titration of mutational burden.

Beyond modeling cancer, our data strongly indicate that our system expands AT2 organoids recapitulating various critical aspects of normal AT2 biology. Notably, we describe culture conditions that induce full and partial differentiation into the AT1 state, which can serve as a powerful model to study the molecular processes governing differentiation and regeneration. Given the feasibility of genetic engineering and comprehensive molecular profiling at the single-cell level, these processes can be dissected in a high-throughput manner with chemical or genetic screens that use sci-ATAC-seq or single cell RNA-seq as a readout (e.g. Perturb-ATAC^51^). Similar approaches could be applied to any other aspect of AT2 biology using our system.

The observation that 20% of organoid lines are not phenotypically stable is not surprising given similar observations in first generation and current organoid platforms^15,16^ as well as current systems^19^. Given that the starting population consists of sorted AT2 cells, we believe that a subset of these cells could be trans-differentiating into pulmonary basal cells. Future studies could shed light on what causes this phenotypic shift, which might be relevant in diseases such as idiopathic pulmonary fibrosis^52,53^. Nevertheless, the majority of our lines (∼80%) maintain their alveolar identity for the entirety of their propagation (∼8 passages). Lastly, for practical usage of our system and to allow organoid derivation from mice with any desired genotype, we identify surface markers (MHCII, EGFR) to rapidly assess phenotypic stability over time.

Our organoid culture system significantly improves on first generation and more recent platforms. In contrast to the protocols described by *Youk et al*.^19^ and *Salahudeen et al*.^18^, the organoids described here expanded rapidly (6-12 day expansion period) and maintained their alveolar identity for at least 8 passages. Rapid expansion and cell-state stability are indispensable features that enabled us to model complex genotypes and various environmental perturbations. The system developed by *Katsura et al*.^20^ is similarly fast (10-14 day expansion period) and maintains AT2 marker expression for at least 6 passages. Our media formulation differs from *Katsura et al*.’s in two major ways—while we use HFG to stimulate the c-MET pathway, they employ IL-1β and EGF to activate inflammatory signaling and the EGFR cascade, respectively. Therefore, our platform provides an equally powerful yet distinct way to support alveolar organoid growth compared to the current state-of-the-art technology.

Given the significant advances of our culture system and its flexibility to rapidly and faithfully model lung cancers of diverse genotypes, we envision transformative applications in normal alveolar stem cell biology and tumor evolution.

## MATERIALS AND METHODS

### Tissue processing for organoid culture

8-17 week old mice were sacrificed and their lungs were inflated with digestion buffer containing Advanced DMEM/F-12, Penicillin/Streptomycin, Amphotericin B, 1 mg/mL Collagenase (Sigma, C9407-500MG), 40 U/mL DNase I (Roche, 10104159001) 5µM HEPES, and 0.36 mM CaCl_2_. Distal lung tissue was extracted, minced and incubated in 3-5mL digestion buffer at 37°C for 30-45 min with agitation. The resulting suspension was washed two times with 1x PBS, filtered through 70µm mesh to remove chunks, and incubated in ACK Lysis Buffer (Thermo, A1049201) for 3-5 minutes at room temperature to lyse red blood cells. The suspension was washed two more times and resuspended in 1x PBS and processed further for sorting (see below).

### Organoid culture

AT2 cells were sorted from dissociated lung cell suspensions as previously described^35^. Briefly, cells were resuspended in FACS buffer containing 1x PBS, 0.1% BSA, and 2 mM EDTA then stained with anti-mouse CD31-APC (1:500, Biolegend, 102507), CD45-APC (1:500, BD Biosciences, 559864), EpCAM-PE (1:500, Biolegend, 118206), MHCII-APC-eFluor-780 (1:500, Thermo, 47-5321-82) on ice for 30 minutes and then resuspended in FACS buffer containing DAPI (1µg/mL, Thermo, D1306). The suspensions were then sorted for DAPI-, CD31-, CD45-, EpCAM+, MHCII+ cells. Approximately, 10,000 sorted AT2 cells were mixed with Growth Factor Reduced Matrigel (Corning) at a ratio of 1:9 and seeded onto multi-well plates as 20µL drops. The drops were incubated at 37°C for 15 minutes – 3 hours to allow them solidify, then overlaid with F^7^NHCAS or F^7^NHCS medium supplemented with Y-27632 (see below).

The cultures were maintained in a humidified 37°C / 5% CO_2_incubator at ambient O_2_ pressure. Media was replenished every 3-4 days using F^7^NHCAS or F^7^NHCS medium without Y-27632 and organoids were passaged 6-12 days after plating. For passaging, matrigel drops were dissolved in TrypLE Express (Sigma, 12604-013) and incubated at 37°C for 7-15 minutes. The organoid suspensions were then dissociated into single cells by vigorous pipetting, washed twice, resuspended in 1x PBS, and plated as described above. We typically plated 10,000 cells per drop.

### Organoid medium recipe

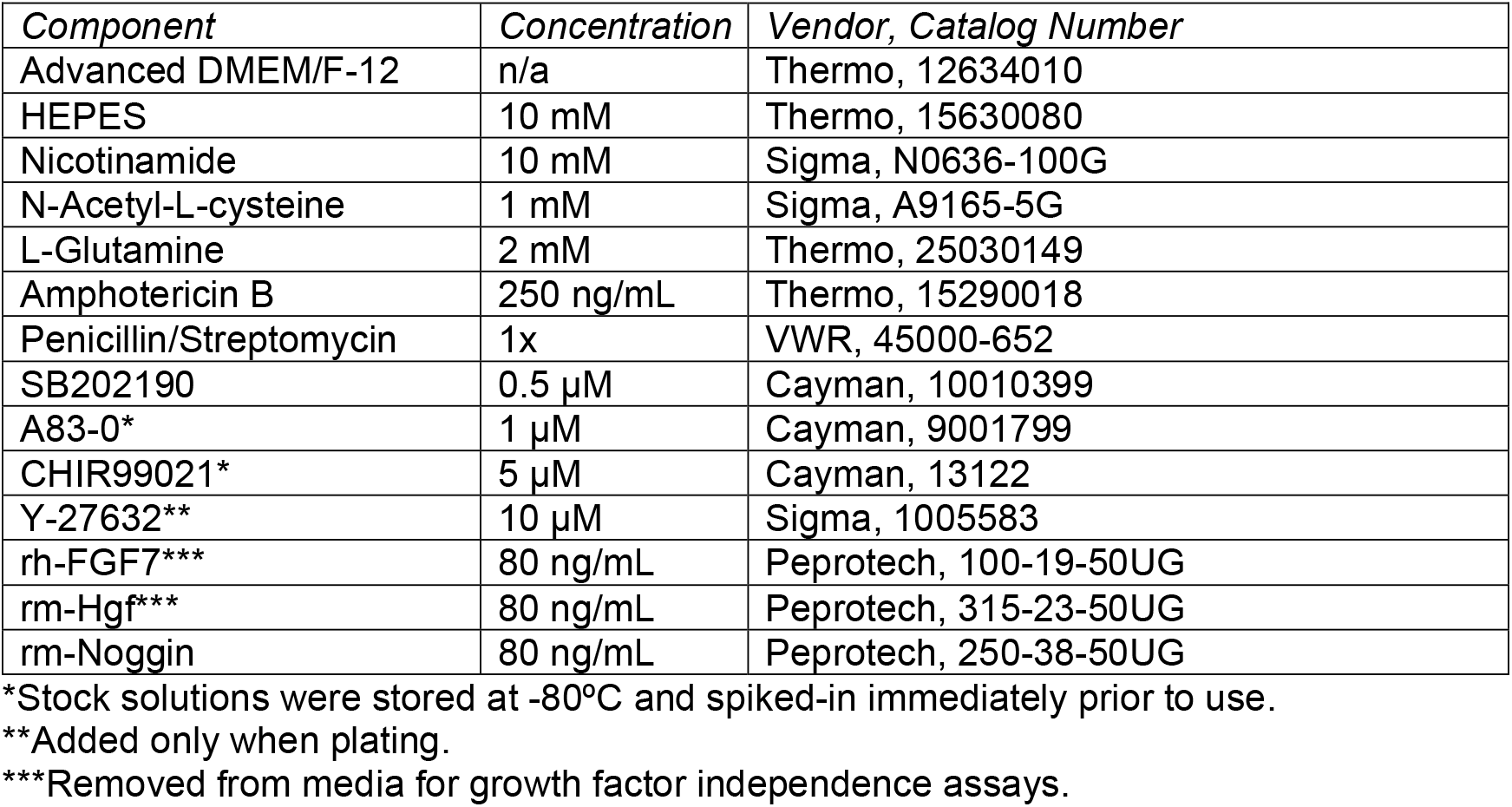

### Flow cytometric analysis

For longitudinal tracking of Sftpc expression, the percentage of cells expressing eGFP was determined via flow cytometry at each passage. For staining cell-type specific surface markers on murine organoids, cells were incubated with either anti-mouse MHC Class II APC-eFluor 780 (1:1000, Thermo, 47-5321-82) or Egf Alexa Fluor 647 (200 ng/mL, Thermo, E35351) antibodies on ice for 20 minutes. Flow cytometric analysis was performed on a Guava EasyCyte flow cytometer. Data analysis was performed using FlowJo software.

### Organoid Viral transduction

Viral transduction of organoids was carried out using the previously described “mix-and-seed” method^54^. Briefly, organoids were processed for passaging, resuspended in concentrated virus, mixed with matrigel, and plated. We do not recommend using spinfection-based protocols described in the literature because the cells tend to attach to most surfaces, even those that have not been treated for tissue culture. Adenoviral transductions were performed at an MOI of 250-300. Lentiviral transductions were performed using non-tittered viruses. The Ad5-sgEA-Cbh-Cre virus was purchased from ViraQuest Inc. The Ad5-CMV-Cre virus was purchased from the Viral Vector Core at University of Iowa Carver College of Medicine.

### Animal studies

Mice were housed at the animal facility at the Koch Institute for Integrative Cancer Research at MIT. All animal studies described in this study were approved by the MIT Institutional Animal Care and Use Committee. *Kras*^*LSL-G12D/+* 1^, *Trp53*^*fl/fl* 55^, *Rosa26*^*LSL- Cas9-eGFP* 56^, *Sftpc-eGFP* ^30^ mice have been previously described and were maintained on a pure C57BL/6 background.

### Organoid transplants

For all transplant experiments, mice were intratracheally inoculated with 40 µL bleomycin (0.1 mg/kg, Cayman, 13877) and organoids were seeded on the same day. Whole organoids were harvested 3 days later using dispase (Corning, 354235) and approximately 1×10^4^ whole organoids were transplanted orthotopically into syngeneic recipients via intratracheal delivery. An approximately even distribution of male and female transplant recipients at 8-20 weeks of age were used for all experiments.

### Tumor analysis

Mice receiving *KP* and *EAP* organoids were sacrificed 16 and 8 weeks post-transplantation, respectively. Lungs were immediately scrutinized for green fluorescent nodules using a dissecting microscope. A positive score was assigned to mice bearing at least one fluorescent lesion. Histopathological analysis was conducted using a deep neural network developed by Aiforia Technologies and the Jacks lab, with consultation from the veterinarian pathologist, Dr. Roderick Bronson^38,57^.

### Histology

Organoids were gently dissociated with wide bore pipet tip, washed in 1xPBS, and pelleted at 100 x g for 5 min. Organoid pellets were resuspended gently in 4% PFA for 2 hours at 4C, transferred to 70% EtOH, and embedded in paraffin. Tissues were fixed in zinc formalin (Polysciences, 21516) overnight at room temperature and maintained in 70% EtOH before being processed for paraffin embedding.

### Immunohistochemistry (IHC)

Sectioned organoids or tissues were stained with hematoxylin and eosin (H&E) or IHC stained with the following antibodies: *organoids*: anti-TTF1 (1:500, Abcam, ab76013), rabbit anti-SFTPC (1:5000, Millipore, ABC99), rabbit anti-Caveolin-1 (1:5000, Thermo, C3237), rabbit anti-CCSP (1:5000, Millipore, 07-623), rabbit anti-Keratin 5 (1:5000, Biolegend, 905501), anti-TTF1 (1:500, Abcam, ab76013); *tissues*: rabbit anti-TTF1 (1:500, Abcam, ab76013), rabbit anti-HMGA2 (1:400, Cell Signaling Technology, #8179). For IHC staining of lung and organoid sections, antigen retrieval was performed in citrate buffer (10 mM, pH 6.0) at 97 °C for 20 min. Slides were processed using a Thermo Scientific Autostainer 360 with the following run conditions: endogenous peroxidases blocking for 10 min, protein block for 30 minutes, primary antibody for 60 minutes, labelled polymer for 30 minutes.

### Fresh frozen lung preparation and immunofluorescence microscopy

Lungs were harvested from mice and a 50% OCT media/ 50% PBS mixture was injected into the lungs via the trachea. Lungs were then placed in a cryomold and covered in 100% OCT media. Cryomolds were floated on a solution of 2-methylbutane cooled by dry ice. 7 μm sections were cut on a cryostat and sections were stored at – 20. The day of staining, sections were fixed in 4% paraformaldehyde for 5 minutes at room temperature. Sections were then washed 3 times with PBS, and then blocked with 5% bovine serum albumin in PBS for 1 hour at room temperature. Sections were stained with rabbit anti-SFTPC (1:400, Millipore, ABC99), rabbit anti-Caveolin-1 (1:400, Thermo, C3237), anti-CD45 (1:200, Biolegend, 30-F11). Donkey anti-rabbit secondary antibodies were used to detect SFTPC and Caveolin-1 antibodies (1:600, Jackson Immunoresearch). Sections were washed and mounted with Prolong Diamond with DAPI (Thermo Fisher Scientific). Images were acquired using a Nikon Eclipse 80i epi-fluorescent microscope.

### Bulk transcriptome analysis

Total RNA was isolated from sorted cells using the TRIzol Plus RNA Purification Kit (Invitrogen, 12183555). Briefly, cells were sorted directly into TRIzol. Following lysis and phase separation, total RNA was purified from the aqueous phase using the PureLink RNA Mini Kit (included in the TRIzol Plus RNA Purification Kit) according to the manufacturer’s specifications. RNAseq libraries were prepared from 300ng of total RNA using the Kapa Hyperprep kit (Roche) with 14 cycles of PCR. RNAseq libraries were quality controlled using Fragment Analysis (Agilent) and qPCR and pooled for sequencing. The libraries were sequenced with single-end 75 base pair reads on an Illumina NextSeq Instrument. Sequence reads were trimmed to eliminate 3’ adapter traces using cutadapt (v1.16)^58^. Trimmed reads were aligned to the mouse genome (mm9 build, UCSC annotation, genome.ucsc.edu) with STAR (v2.5.3a)^59^. Reads per feature were quantified using the featureCounts utility in the Subread package (v1.6.2)^60^. Differential analyses were performed using DESeq2 (v1.28.1)^61^ on raw counts. Enrichment analyses were performed using GSEA (v4.0.3) in the “pre-ranked” mode using DESeq2 reported log2 fold-change values as the ranking metric. The c2 (curated) collection from MSigDB (www.gsea-msigdb.org/gsea/msigdb) was used with the customized addition of four gene-sets (AT2 markers from ^33^; AT1 markers from ^33^; Club markers from ^33^; Pulmonary basal cell markers from ^34^).

### Processing and analysis for sci-ATAC-seq

Cells were prepared for sciATAC-seq as previously described^38,62^.

#### Sci-ATACseq counts generation and QC

Using the generated peak list, the number of reads for each peak window were determined for each barcode tag. This generated a matrix which associated ATAC reads in peaks for each single cell. Only cells with FRIP ≥ 0.3 and 1000 unique nuclear peaks were retained for downstream analysis. After this quality check, 2,444 single cells were retained for further analysis.

#### Single-cell visualization

The matrix of k-mer accessibility deviation Z-scores was first column-scaled and centered (using the scale function in R v.3.5.3) (R Core Team, 2019), and run through a principal component analysis (PCA) dimensionality reduction. The Uniform Manifold Approximation and Projection (UMAP) algorithm (McInnes et al., 2018) was then applied to project single cells in two dimensions using the k-mer PC scores for the first 20 PCs (implemented using the uwot package (v0.1.4) in R with the following non-default clustering parameters: n_neighbors = 20, min_dist = 0.4, metric = ‘‘cosine’’).

#### Gene scoring

Single-cell gene scores were determined as previously described (LaFave et al. 2020). Scores were then normalized to the mean gene score per cell for further use in downstream analysis. For visualization of the gene scores in single cells, the gene scores were smoothed based on their nearest-neighbors (k=30) in PC space. To assess correlation of AT2, AT1, basal, and club signatures to each individual single-cells, AT1 and AT2 signatures were used from normal single-cell ATAC-sequencing data previously described^38^ or from published signatures^33,34^. Gene listed were intersected with the gene score matrix. Signature genes were mean normalized and Z scores were visualized on the UMAP.

### Primer and sgRNA sequences

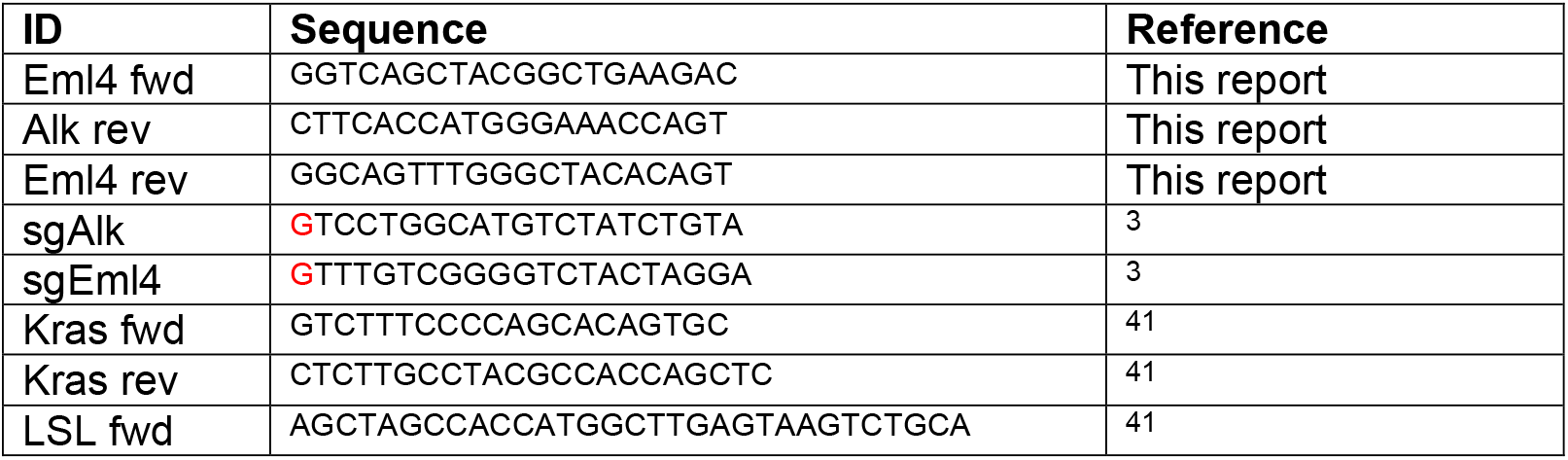

## FIGURE LEGENDS

**Figure S1.**
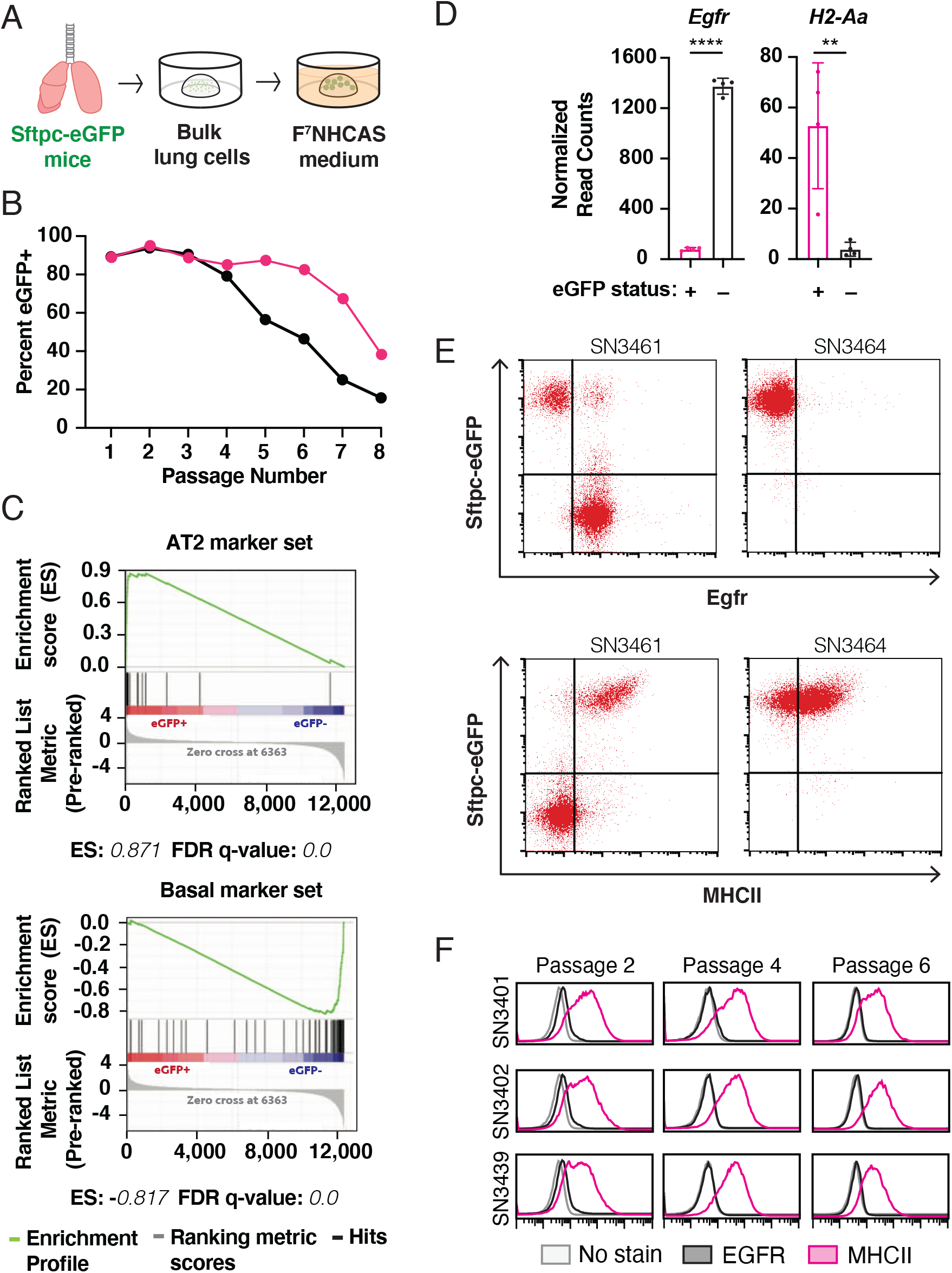
A. Preliminary method for culturing alveolar organoids from normal lungs. B. Flow cytometry-based quantification of Sftpc-eGFP expressing cells over time in organoid culture. A total of *n* = 2 independent lines were established from Sftpc-eGFP mice. C. GSEA enrichment plots of AT2 and basal cell signatures comparing Sftpc-eGFP positive and negative organoids. ES and FDR q-values are shown beneath their corresponding plots. D. *Egfr* and *H2-Aa* (a component of MHCII) expression by RNA sequencing in Sftpc-eGFP positive (pink) and negative (black) organoids. Data are expressed as mean values ± the standard deviation. Statistical analysis in was performed using a two-tailed students t-test. ^****^ denotes a P-value < 0.0001 and ^**^ indicates a P-value = 0.0079. E. Surface expression of EGFR (top) and MHCII (bottom) in Sftpc-eGFP expressing organoids in representative phenotypically unstable (SN3461) or stable (SN3464) lines by flow cytometry. F. Surface expression of EGFR (black) and MHCII (pink) in three independent organoid lines at various passages by flow cytometry.

**Figure S2.**
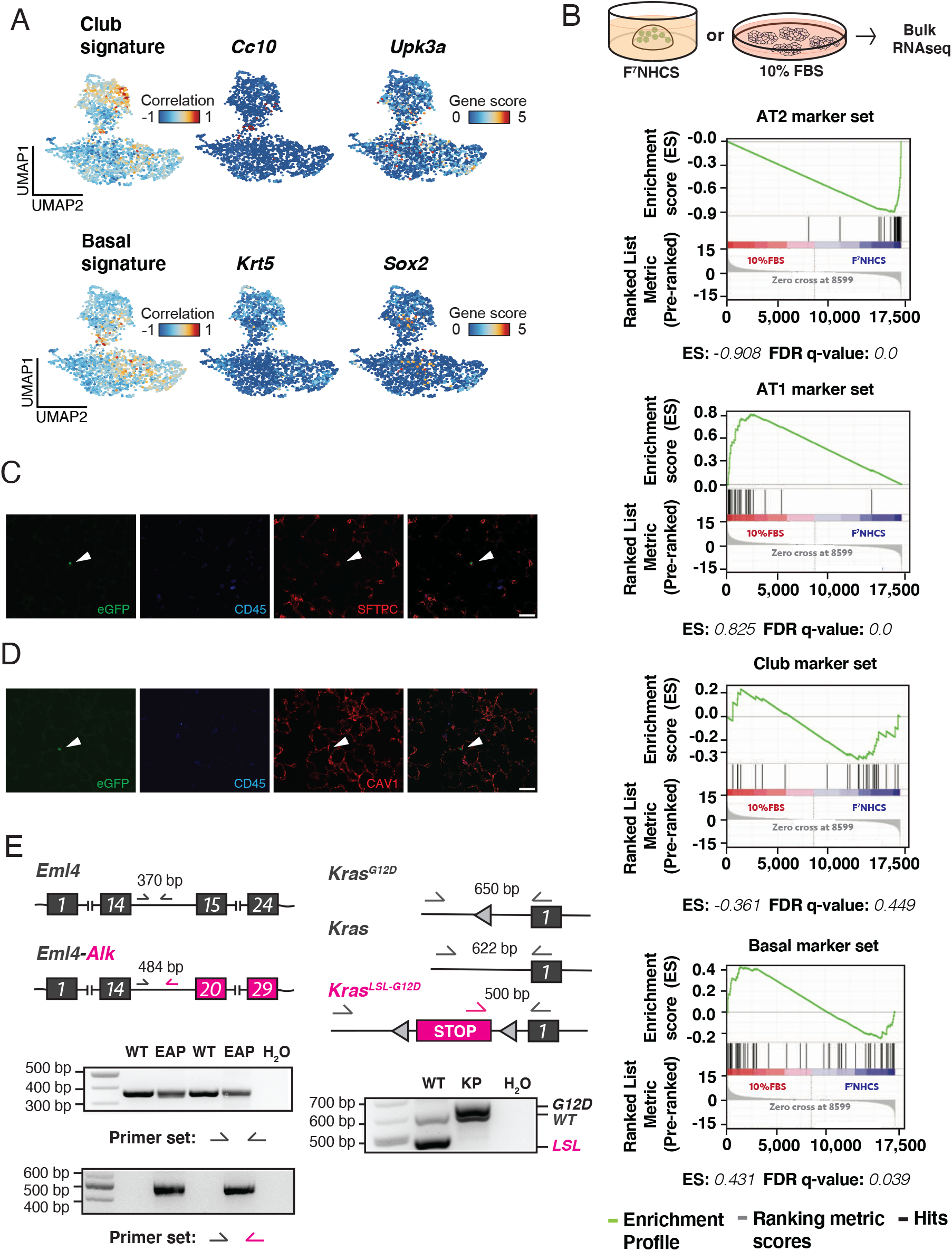
A. Correlation of chromatin profiles in single organoid cells to established gene sets and two individual canonical markers for Club (top) and Basal (bottom) cells. Gene set plots are colored by their Pearson’s differential correlation coefficients. Marker plot color intensity indicates the relative gene score. B. (Top) Schematic for *in vitro* differentiation experiment and (bottom) GSEA enrichment plots of AT2, AT1, Club and Basal marker gene sets comparing organoids cultured in differentiation (10% FBS) or complete AT2 (F^7^NHCS) media. ES and FDR q-values are shown beneath their corresponding plots. C. Immunofluorescence staining of bleomycin preconditioned lung sections 14 days post intra-tracheal transplantation of eGFP-labeled AT2 organoids for eGFP, CD45, and SFTPC. Arrows point to engrafted cells. D. Immunofluorescence staining of bleomycin preconditioned lung sections 14 days post intra-tracheal transplantation of eGFP-labeled AT2 organoids for eGFP, CD45, and CAV1. Arrows point to engrafted cells. E. PCR-based detection of oncogenic mutations in *Alk* and *Kras*. The diagram on the left shows *Eml4* locus before (top) and after the inversion (bottom). The diagram on the right illustrates the *WT* Kras allele (middle) and the *Kras*^*LSL-G12D*^ allele before (bottom) and after (top) recombination. PCR reactions were carried out on genomic DNA isolated from two pairs of *WT* (parental) and *EAP* (mutated) organoids as well as one pair of *WT* (parental) and *KP* organoids using the indicated primer pairs. Expected product sizes are indicated for each primer set.

## ACKNOWLEDGEMENTS AND DISCLOSURES

We thank Carla Concepcion, Sheng Rong Ng, Tuomas Tammela, Francisco J. Sánchez-Rivera, Grissel Jaramillo, Alex Jaeger, Caterina Colon, Demi Sandel, William Rideout III, Kim Mercer, Megan Burger, William Freed-Pastor and the rest of the extended Jacks Lab family for helpful discussions and technical assistance; George Eng, Jonathan Braverman, and Omer Yilmaz for helpful advice regarding organoid culture; The MIT BioMicro Center for performing high-throughput sequencing; the Koch Institute’s Robert A. Swanson (1969) Biotechnology Center for technical support, specifically the Histology Core Facility, the Flow Cytometry Core Facility and the Nanotechnology Materials Lab; Karen Yee and Judy Teixeira for administrative support; and Quincy Phillips and Hildy Grossman for supporting this work.

This work was supported by the Howard Hughes Medical Institute, the Koch Institute Support Grant P30-CA14051 from the National Cancer Institute, and the Koch Institute Frontier Research Program through gifts from Upstage Lung Cancer. S.N. was supported by Howard Hughes Medical Institute Gilliam Fellowship Program and the David H. Koch Graduate Fellowship Fund. S.N., C.C., and R.R. were supported in part by the NIH Pre-Doctoral Training Grant T32GM007287.

T.J. is a member of the Board of Directors of Amgen and Thermo Fisher Scientific. He is also a co-Founder of Dragonfly Therapeutics and T2 Biosystems. T.J. serves on the Scientific Advisory Board of Dragonfly Therapeutics, SQZ Biotech, and Skyhawk Therapeutics. He is the President of Break Through Cancer. None of these affiliations represent a conflict of interest with respect to the design or execution of this study or interpretation of data presented in this manuscript. T.J. laboratory currently also receives funding from the Johnson & Johnson Lung Cancer Initiative and The Lustgarten Foundation for Pancreatic Cancer Research, but this funding did not support the research described in this manuscript.

## Notes

### Competing Interest Statement

The authors have declared no competing interest.

